# The Relationship between Parasitic Fleas and Small Mammals in Household of Western Yunnan Province, China

**DOI:** 10.1101/518266

**Authors:** Jia-Xiang Yin, Xiao-Ou Cheng, Qiu-Fang Zhao, Zhao-Fei Wei, Dan-Dan Xu, Meng-Di Wang, Yun Zhou, Xiu-Fang Wang, Zheng-Xiang Liu

## Abstract

The Yunnan Province has one of the most serious outbreaks of the plague epidemic in China. Small mammals and fleas are risk factors for the occurrence of plague in commensal plague foci. Understanding the relationship between parasitic fleas and small mammals will help control fleas and prevent the onset of the plague. Four hundred and twenty-one small mammals, belonging to 9 species, were captured. Of these, 170 small mammals (40.4%) were infested with fleas. A total of 992 parasitic fleas (including 5 species) was collected. The number of *Leptopsylla Segnis* and *Xenopsylla Cheopis* was 91.0%. The final multiple hurdle negative binomial regression model showed that when compared with *Rattus Tanezumi*, the probability of flea infestation on *Mus musculus* and other host species decreased from 58% to 99%, while the infestation with fleas from other host species increased 4.7 fold. The probability of flea prevalence in adult hosts increased by 74%, while the number of fleas decreased by 76%. The number of flea infestations in small male mammals increased by 62%. The number of fleas in small mammals weighing more than 59 grams has been multiplied by about 4. *Rattus Tanezumi* is the predominant species in households in West Yunnan Province, while *Leptopsylla Segnis* and *Xenopsylla Cheopis* are dominant parasitic fleas. There is a strong relationship between the abundance of parasitic fleas and the characteristics of small mammals (e.g. Species, age, sex, and body weight).

## Introduction

Plague is a zoonotic bacterial infectious disease transmitted by fleas with reservoirs in rodent populations, in which the parasitic flea plays a potentially fundamental role as bridge vectors to transmit the bacteria (*Yersinia pestis*) to animals and humans [1]. The potential role of fleas as carriers and host-supported local reservoirs can help explain the persistence of plague [2].

The plague has played a huge role in human history and is still prevalent in most parts of China. In China, animal plague is reported almost every year, and occasional human plagues occur [3, 4]. Transmission to humans sometimes occurs through contact with fleas that have fed on an infected small mammal or by skinning infected small mammals (e.g. Marmot) [5–7]. The province of Yunnan is one of the most serious plague epidemic foci in China [8,9], and especially the most active in commensal rodent plague foci [2]. The province of Yunnan is characterized by complex natural conditions, an obvious vertical climate, consisting of embedded microhabitats that create a series of ecological niches, which may increase the risk of establishing vectors and specific hosts that carry pathogens.

Fleas are obligatory hematophagous insects and rodents are the most common host of fleas, although they can appear in other types of mammals and birds [10, 11]. At different stages of the life cycle, fleas are located in the hosts or in the host's nest. In most cases, the imago flea is usually parasitic on the host and until the end of its life cycle, differentiating with the larvae. When leaving a pupa, a new flea of imago finds a host because the reproduction of the flea is not possible without the feeding of blood [12]. Therefore, it is likely to handle flea infestations on live hosts. The parasites get their food and other biological supplies from their hosts. Consequently, they can affect the host directly by reducing host resources, and indirectly by provoking behavioral responses [13, 14] or immunological responses [1, 15]. Fleas with high density and richness not only favor the bacteria plague, but also spread widely among the hosts, but also suppress the immune response of the hosts, thus improving the susceptibility of the hosts [16, 17], which is vital for the maintenance of the plague [18, 19].

The abundance of hosts is an important factor that affects the distribution and abundance of parasites. In terms of the reservoir, some individuals, populations or host species are characterized by a higher level of parasite infestation than others [20]. It is well known that there is a great variation in the abundance of a parasite among the different host species. The host with the highest abundance of parasites is commonly defined as the main host, while those with lower abundance of parasites are defined as auxiliary hosts [20, 21].

In this study, we attempt to describe the distribution of small parasitic mammals and fleas and reveal the effect of small mammals on the abundance of parasites in homes in the western province of Yunnan. The information on the factors that affect the flea abundance in hosts is essential to provide evidence-based recommendations on flea control and to implement plague control and prevention programs in natural plague foci.

## Methods

### Description of the study area

The western of Yunnan province is located in the central region of the Hengduan mountains of the terrestrial part of the Tibetan plateau, which belongs to the low latitude mountain valley. It is influenced by the enormous difference in latitude and altitude gradient, which is characterized by complex terrain and varied climate [22]. Therefore, it is one of the regions that possess the most abundant animal and plant species in China. There are many minority nationalities, including Yi, Dai, Naxi, Hui, Bai et al. inhabiting this area. The main economic source is planting and economic development is relatively poor. These conditions contribute to the breeding of small mammals and the spread of the plague.

During the period from July 2011 to October 2012, the study was carried out in 800 households in 40 villages, 10 counties, in the western part of Yunnan province (Fig 1). In the study area, the 10 counties have a total area of approximately 41,130 square kilometers and a population of approximately 3.4 million.

**Figure 1.**
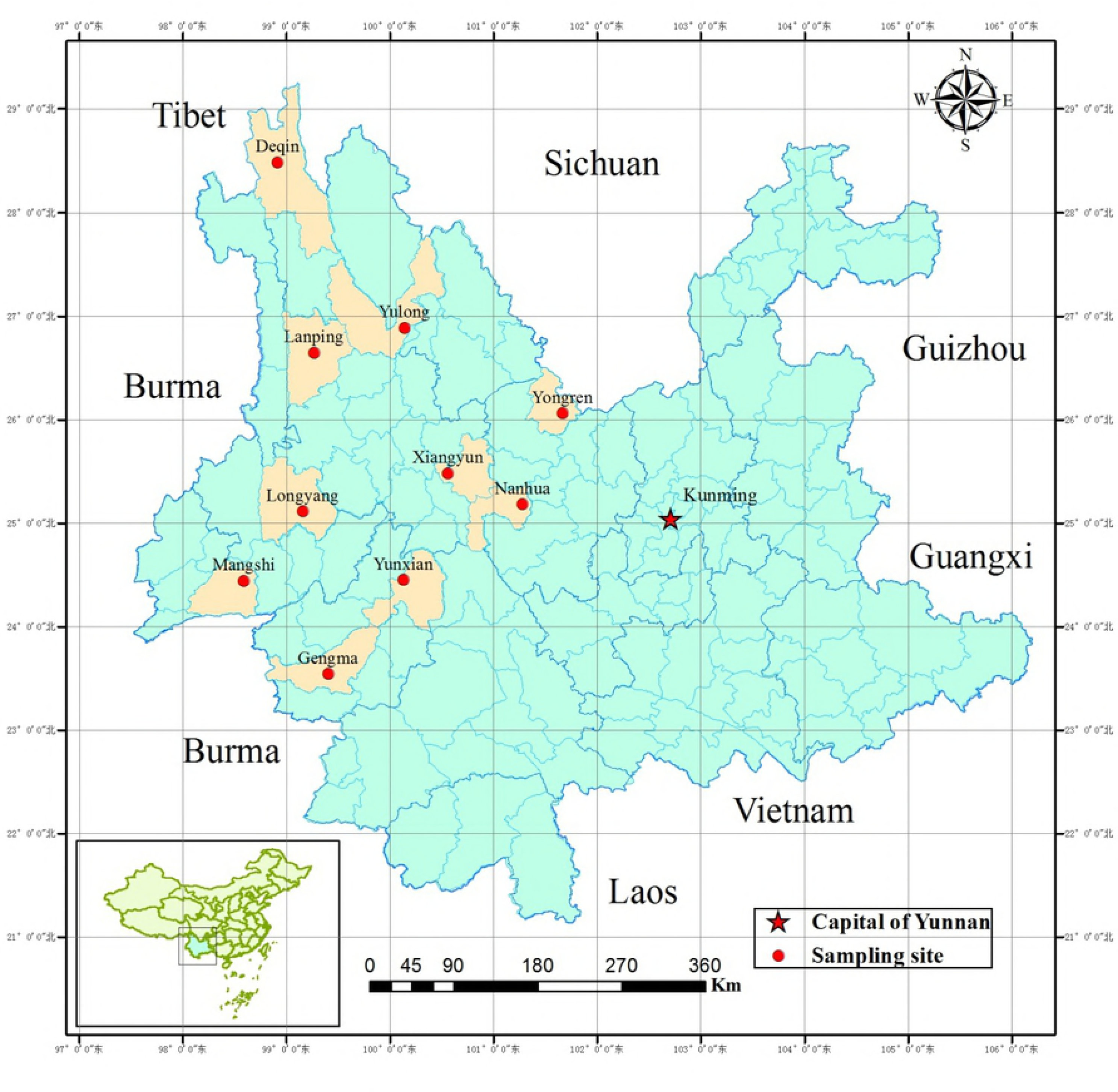
Location of 10 sampling sites in Western Yunnan Province, China. Ten red dots represent 10 sampling counties and yellow shadow is the acreage of each county. Red five-point star is the location of Kunming city (capital of Yunnan Province).

### Small mammals capture and identification

Five live traps were placed in each home for three continuous nights to capture small mammals. The bait was fried ham sausage. Each live trap was verified on the morning of the next day. If a small mammal was caught, a new live trap with fresh bait was replaced by the old live trap and placed in the same place. The species of small mammals, sex and age were identified according to their morphological characteristics. Their weight, body length and tail length were measured.

### Parasitic flea collections and identification

The parasitic fleas were harvested by combing from the tail end forward using a toothbrush in a white enamel tray after anesthetizing small mammals with ether. The parasitic fleas were stored in labeled vials containing 75% ethanol. Fleas from each small mammal were kept in separate vials and kept at room temperature. Flea species were identified under a light microscope.

### Data analysis

The distribution of small parasitic mammals and fleas was summarized using descriptive statistics. Then the abundance of parasitic fleas of each small mammal was generated. Species, sex, age, weight, body length and tail length of each small mammal were considered potential factors for the abundance of parasitic fleas. For continuous variables, the median was used to classify these variables into binary variables. The relationship between potential factors and a result was explored using the hurdle binomial regression model (HNB) of the regression model under R software 3.02 [23–25]. This regression model was applied to take into account the current data set that shows a count of excess zeros and an overdispersion distribution. It is a two-component model: one is the logistic model adjusting counts against zero, the other is the hurdle binomial model adjusting positive counts. Potential factors (*P*<0.20) related to small mammals entered the prototype multiple regression model HNB. The final model to predict the factors associated with the parasitic flea abundance was refined using a backward method (α = 0.10 as a criterion of statistical significance). The proportion of Prevalence odds ratio (OR) and abundance ratio (AR) for a parasitic infestation of fleas were calculated based on pieces of literature [26,27].

## Results

### Distribution and description of small mammals

A total of 12,000 traps was placed in 800 households in 40 villages in 10 counties in western Yunnan Province and 421 small mammals were captured. The overall density of small mammals was 3.51%. Small mammals were divided into 9 species, 6 genera, 2 families and 2 orders (Table 1). *Rattus Tanezumi* is the dominant species and it represents 66.03% (278/421) of the entire sample. Among the 10 counties, the highest number and the smallest number of small mammals were captured in Mangshi (100) and Deqin (16) counties, respectively. The number of small mammals richness was 9. The highest number and lowest number of richness were in Yongren (5), Nanhua (1) and Deqin (1) counties, respectively.

**Table 1.** Distribution for small mammals in households of 10 counties in Western Yunnan Province.

| County | Small mammals |  |  | Species and proportion (%) of small mammals |  |  |  |  |  |  |  |  |
| --- | --- | --- | --- | --- | --- | --- | --- | --- | --- | --- | --- | --- |
|  | Abundance | Density | Richness | *Rattus* | *Mus* | *Rattus* | *Rattus* | *Suncus* | *Apodemus* | *Crocidura* | *Anoro sorex* | *Mus* |
|  |  |  |  | *Tanezumi* | *musculus* | *norvegicus* | *Nitidus* | *Murinus* | *Chevrieri* | *Attenuata* | *squamipes* | *caroli* |
| Yongren | 86 | 7.17 | 5 | 4(4.65) | 73(84.88) | 2(2.33) | 6(6.98) | 0 | 0 | 0 | 0 | 1(1.16) |
| Nanhua | 36 | 3.00 | 1 | 36(100.00) | 0 | 0 | 0 | 0 | 0 | 0 | 0 | 0 |
| Xiangyun | 41 | 3.42 | 3 | 18(43.90) | 0 | 22(53.66) | 0 | 1(2.44) | 0 | 0 | 0 | 0 |
| Yunxian | 39 | 3.25 | 2 | 37(94.87) | 2(5.13) | 0 | 0 | 0 | 0 | 0 | 0 | 0 |
| Gengma | 26 | 2.17 | 2 | 25(96.15) | 1(3.85) | 0 | 0 | 0 | 0 | 0 | 0 | 0 |
| Mangshi | 100 | 8.33 | 2 | 94(94.00) | 0 | 0 | 0 | 6(6.00) | 0 | 0 | 0 | 0 |
| Longyang | 28 | 2.33 | 3 | 24(85.71) | 0 | 0 | 0 | 0 | 0 | 2(7.14) | 2(7.14) | 0 |
| Lanping | 22 | 1.83 | 3 | 17(77.27) | 0 | 4(18.18) | 1(4.55) | 0 | 0 | 0 | 0 | 0 |
| Yulong | 27 | 2.25 | 3 | 23(85.18) | 0 | 2(7.41)) | 0 | 0 | 2(7.41) | 0 | 0 | 0 |
| Deqin | 16 | 1.33 | 1 | 0 | 0 | 0 | 16(100.00) | 0 | 0 | 0 | 0 | 0 |
| Total | 421 | 3.51 | 9 | 278(66.03) | 76(18.05) | 30(7.13) | 23(5.46) | 7(1.66) | 2(0.48) | 2(0.48) | 2(0.48) | 1(0.24) |

\*Small mammals density = (abundance/ live traps.nights)×100%, the number of live traps.nights each county is 1200

In our study, the weight, body length and tail length of 407 small mammals were calculated (for other 14 small mammals there was no calculation due to incomplete data). The mean body weight was 62.93g, standard deviation 42.67g, median 59.31g (range: 7.39-186.89g); the average body length was 22.79cm, the standard deviation of 8.57cm and the median 21.00cm (range: 7.50 to 39.30cm); the average length of the tail was 12.30cm, the standard deviation of 3.76cm and the median 13.30cm (extremes: 1.00 and 20.00cm).

### Distribution of parasitic fleas

Of 421 small mammals, 170 small mammals were infested with fleas and their prevalence was 40.38%. A total of 992 parasitic fleas, divided into 5 species, 5 genera, and 3 families, were collected and the flea value was 2.36 (992/421). The highest prevalence of fleas and the highest flea index occurred in Yunxian County (71.79%) and Yulong County (7.33), respectively. Out of 992 fleas, the number of *Xenopsylla cheopis* and *Leptopsylla segnis* was 903, or 91.03%. The proportion of *Xenopsylla cheopis* exceeds 90% in Yunxian and Gengma counties. (Table 2)

**Table 2.** Distribution for parasitic fleas in households of 10 counties from Western Yunnan Province.

| County | No. of small mammals | No. of small mammals with flea infection | Flea prevalence (%)<sup>a</sup> | No. of fleas | Flea index<sup>b</sup> | Richness | Parasitic flea species and proportions |  |  |  |  |
| --- | --- | --- | --- | --- | --- | --- | --- | --- | --- | --- | --- |
|  |  |  |  |  |  |  | *Xenopsylla*<br>*Cheopis* | *Leptopsylla*<br>*Segnis* | *Ctenocephalides*<br>*felis felis* | *Monopychllusani*<br>*sus* | *Pulexirrita*<br>*ns* |
| Yongren | 86 | 6 | 6.98 | 83 | 0.97 | 3 | 71(85.54) | 11(13.26) | 0 | 1(1.20) | 0 |
| Nanhua | 36 | 12 | 33.33 | 86 | 2.39 | 4 | 1(1.16) | 61(70.93) | 8(9.3) | 0 | 16(18.6) |
| Xiangyun | 41 | 21 | 51.22 | 146 | 3.56 | 2 | 3(2.05) | 143(97.95) | 0 | 0 | 0 |
| Yunxian | 39 | 28 | 71.79 | 97 | 2.49 | 2 | 92(94.85) | 5(5.15) | 0 | 0 | 0 |
| Gengma | 26 | 12 | 46.15 | 42 | 1.62 | 2 | 38(90.48) | 0 | 0 | 4(9.52) | 0 |
| Mangshi | 100 | 56 | 56.00 | 250 | 2.50 | 3 | 208(83.20) | 34(13.60) | 0 | 8(3.20) | 0 |
| Longyang | 28 | 11 | 39.29 | 41 | 1.46 | 4 | 16(39.02) | 23(56.09) | 1(2.44) | 1(2.44) | 0 |
| Lanping | 22 | 9 | 40.91 | 49 | 2.23 | 2 | 0 | 45(91.84) | 0 | 4(8.16) | 0 |
| Yulong | 27 | 15 | 55.56 | 198 | 7.33 | 3 | 0 | 152(76.77) | 35(17.68) | 11(5.56) | 0 |
| Deqin | 16 | 0 | 0 | 0 | 0 | 0 | 0 | 0 | 0 | 0 | 0 |
| Total | 421 | 170 | 40.38 | 992 | 2.36 | 5 | 429(43.25) | 474(47.78) | 44(4.44) | 29(2.92) | 16(1.61) |

<sup>a</sup>Flea prevalence = (No. of small mammals with flea infection/ No. of small mammals)×100%
<sup>b</sup>Flea index = No. of fleas/ No. of small mammals

Among the 9 small mammal species, the prevalence of *Rattus Tanezumi* fleas was highest (53.60%, 149/278). The infestation of *Rattus Tanezumi* by *Xenopsylla cheopis* was particularly high (35.61%, 99/278). *Crocidura attenuata* and *Anourosorex squamipes* were not infested with fleas (Table 3).

**Table 3.**
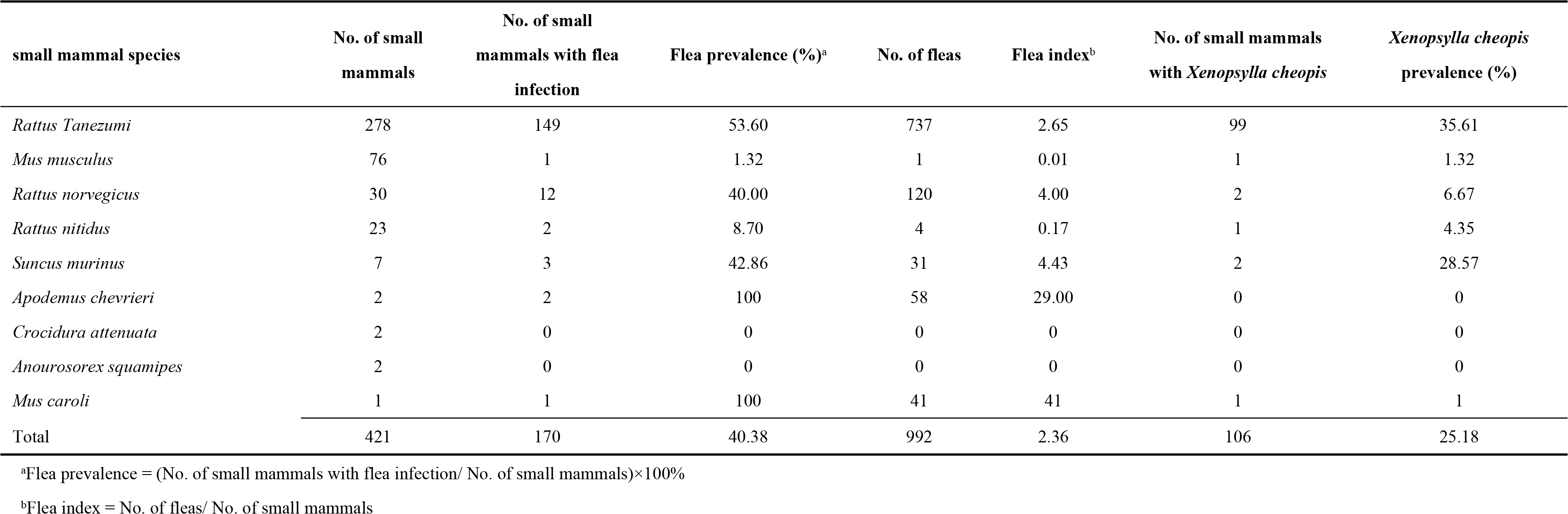
Flea infection for different small mammal species in households of Western Yunnan Province.

| small mammal species | No. of small mammals | No. of small mammals with flea infection | Flea prevalence (%)<sup>a</sup> | No. of fleas | Flea index<sup>b</sup> | No. of small mammals with *Xenopsylla cheopis* | *Xenopsylla cheopis* prevalence (%) |
| --- | --- | --- | --- | --- | --- | --- | --- |
| *Rattus Tanezumi* | 278 | 149 | 53.60 | 737 | 2.65 | 99 | 35.61 |
| *Mus musculus* | 76 | 1 | 1.32 | 1 | 0.01 | 1 | 1.32 |
| *Rattus norvegicus* | 30 | 12 | 40.00 | 120 | 4.00 | 2 | 6.67 |
| *Rattus nitidus* | 23 | 2 | 8.70 | 4 | 0.17 | 1 | 4.35 |
| *Suncus murinus* | 7 | 3 | 42.86 | 31 | 4.43 | 2 | 28.57 |
| *Apodemus chevrieri* | 2 | 2 | 100 | 58 | 29.00 | 0 | 0 |
| *Crocidura attenuata* | 2 | 0 | 0 | 0 | 0 | 0 | 0 |
| *Anourosorex squamipes* | 2 | 0 | 0 | 0 | 0 | 0 | 0 |
| *Mus caroli* | 1 | 1 | 100 | 41 | 41 | 1 | 1 |
| Total | 421 | 170 | 40.38 | 992 | 2.36 | 106 | 25.18 |

<sup>a</sup>Flea prevalence = (No. of small mammals with flea infection/ No. of small mammals)×100%
<sup>b</sup>Flea index = No. of fleas/ No. of small mammals

### The results of the prototype multiple HNB regression model

Table 4 shows the prototype multiple HNB regression model for the abundance of parasitic fleas. Of the six potential factors compared to *Rattus Tanezumi*, the probability of flea infestation of *Mus musculus* and other host species decreased, while the number of flea infestations of other host species increased. The number of fleas in adult hosts decreased; the number of fleas in small mammals weighing more than 59 grams increased.

**Table 4.**
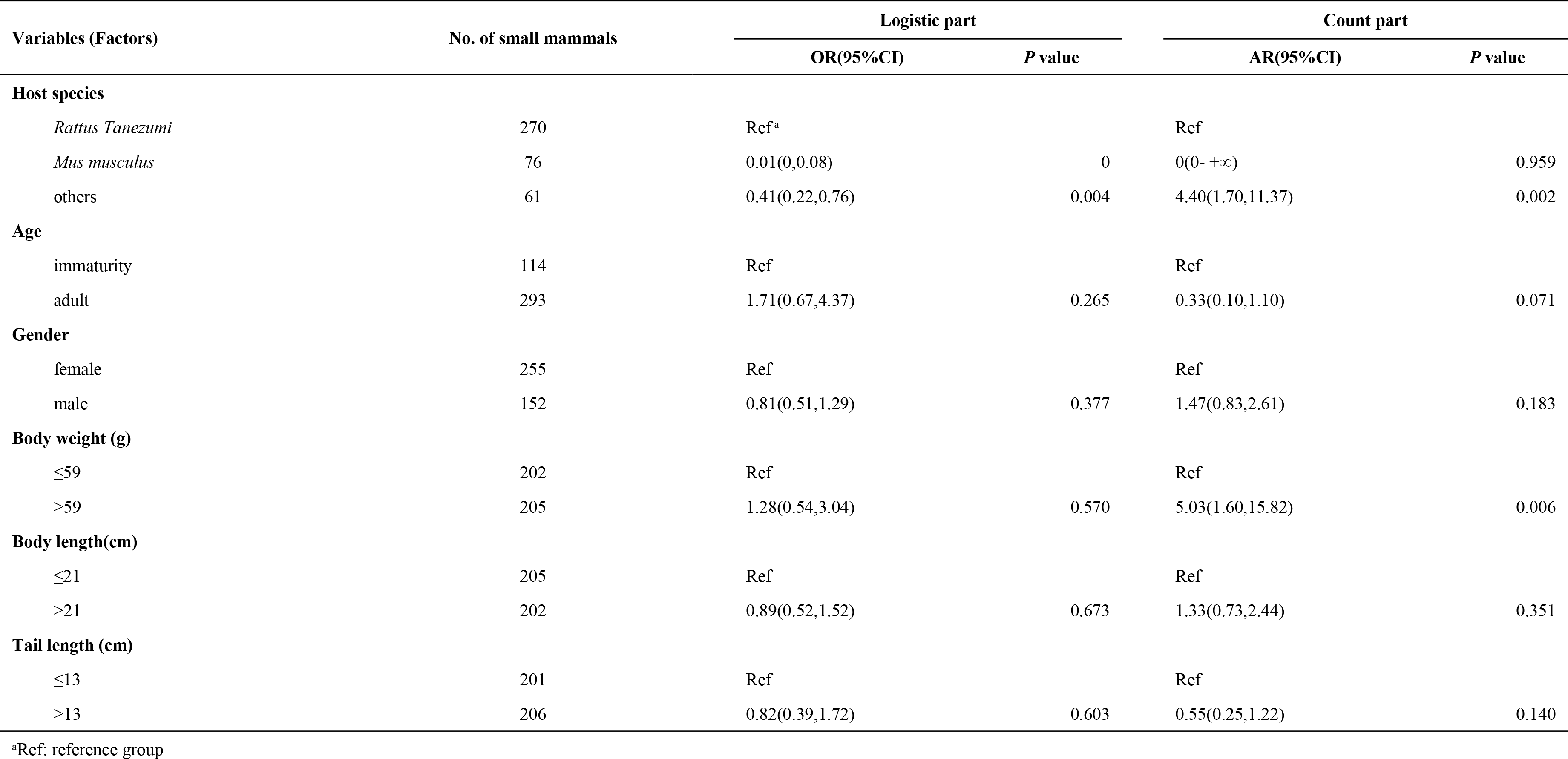
The results of the prototype multiple HNB regression model for the abundance of parasitic fleas.

| Variables (Factors) | No. of small mammals | Logistic part |  | Count part |  |
| --- | --- | --- | --- | --- | --- |
|  |  | OR(95%CI) | P value | AR(95%CI) | P value |
| Host species |  |  |  |  |  |
| *Rattus Tanezumi* | 270 | Ref<sup>a</sup> |  | Ref |  |
| *Mus musculus* | 76 | 0.01(0,0.08) | 0 | 0(0- +∞) | 0.959 |
| others | 61 | 0.41(0.22,0.76) | 0.004 | 4.40(1.70,11.37) | 0.002 |
| Age |  |  |  |  |  |
| immaturity | 114 | Ref |  | Ref |  |
| adult | 293 | 1.71(0.67,4.37) | 0.265 | 0.33(0.10,1.10) | 0.071 |
| Gender |  |  |  |  |  |
| female | 255 | Ref |  | Ref |  |
| male | 152 | 0.81(0.51,1.29) | 0.377 | 1.47(0.83,2.61) | 0.183 |
| Body weight (g) |  |  |  |  |  |
| ≤59 | 202 | Ref |  | Ref |  |
| >59 | 205 | 1.28(0.54,3.04) | 0.570 | 5.03(1.60,15.82) | 0.006 |
| Body length(cm) |  |  |  |  |  |
| ≤21 | 205 | Ref |  | Ref |  |
| >21 | 202 | 0.89(0.52,1.52) | 0.673 | 1.33(0.73,2.44) | 0.351 |
| Tail length (cm) |  |  |  |  |  |
| ≤13 | 201 | Ref |  | Ref |  |
| >13 | 206 | 0.82(0.39,1.72) | 0.603 | 0.55(0.25,1.22) | 0.140 |

<sup>a</sup>Ref: reference group

### The results of the final multiple HNB regression model

The final model showed that four factors - species, age, sex and body weight of small mammals - are closely associated with the abundance of parasitic flea (Table 5). Compared to *Rattus Tanezumi*, the probability of infestation with *Mus musculus* fleas as well as other host species decreased by 58% to 99%, while the number of flea infestations of the other host species increased by 4.71. The probability of flea prevalence in adult hosts increased by 74%, while the number of fleas decreased by 76%. The number of flea infestations in small male mammals increased by 62%. The number of fleas in small mammals weighing more than 59 grams increased by about 4 times.

**Table 5:** The results of the final HNB model for the abundance of parasitic fleas.

| Variables (Factors) | Logistic part |  | Count part |  |
| --- | --- | --- | --- | --- |
|  | OR(95%CI) | P value | AR(95%CI) | P value |
| **Host species** |  |  |  |  |
| *Rattus Tanezumi* | Ref<sup>a</sup> |  | Ref |  |
| *Mus musculus* | 0.01(0,0.07) | 0.000 | 0.00(0- +∞) | 0.956 |
| Others | 0.42(0.23,0.76) | 0.004 | 5.71(2.21,14.77) | 0.000 |
| **Age** |  |  |  |  |
| Immaturity | Ref |  | Ref |  |
| Adult | 1.74(1.07,2.81) | 0.024 | 0.24(0.07,0.80) | 0.020 |
| **Gender** |  |  |  |  |
| Female |  |  | Ref |  |
| Male |  |  | 1.62(0.92,2.84) | 0.093 |
| **Body weight (g)** |  |  |  |  |
| ≤59 |  |  | Ref |  |
| >59 |  |  | 5.03(1.62,15.64) | 0.005 |

<sup>a</sup> Ref: reference group

## Discussion

In this study, 421 small mammals were captured and *Rattus Tanezumi* was the predominant species. A total of 992 parasitic fleas was collected from small mammals. The number of *Leptopsylla segnis* and *Xenopsylla cheopis* was 91.03%. A close relationship between parasitic flea abundance and small mammal traits (including species, age, body weight, sex) was evident in households in Western Yunnan Province.

The results of our investigation showed that *Rattus Tanezumi* is the predominant species in households in western Yunnan Province. *Xenopsylla cheopis* and *Leptopsylla segnis* are the dominant fleas host as well as the dominant fleas host of *Rattus Tanezumi*. These results correspond to the previous study [28]. In the commensal plague foci of the western Yunnan Province, *Rattus Tanezumi* is the main host species, while *Mus musculus* is the auxiliary host species. Yin et al. [26] have suggested that the abundance of parasitic fleas is related to host species. Our data indicated that *Rattus Tanezumi* was more likely to be infested with fleas than *Mus musculus* and other species. The main host may or may not be the species in which the parasite evolved for the first time, but it’s currently home to most individuals in the parasite population [21]. It is therefore believed that *Rattus Tanezumi* is the best species in which the majority of parasites live and the probability of flea infestation on *Mus musculus* as well as other species has decreased compared to *Rattus Tanezumi*. The reason for the increase in the number of fleas in other species could be due to the fact that the number of flea infestations on other species was sufficient, although the number of other host species is lower in other species in our study, like *Apodemus chevrieri*. In addition, the fundamental reasons for the diversity in parasitic abundance between the main host and any auxiliary host are often related to the different successes of exploitation and reproduction of a parasite [29]. Previous studies have shown that the phylogenetic correlation between the main host and the auxiliary host can determine parasitic abundance on its auxiliary hosts, as it should reflect the similarities between host species in terms of physiological, ecological characteristics and immunological [30]. On the other hand, it was found that for flea infestation in small mammals, the abundance of a flea on its auxiliary hosts decreased with increasing phylogenetic distance of these hosts relative to the main host [30]. The mechanisms underlying this model are not yet clear but are supposed to be consistent with the differential performance of a flea on auxiliary hosts, which in turn is correlated with a phylogenetic distance from the auxiliary host to the main host [31].

Recent studies have shown that parasites appear to make favorable choices and decisions for hosts in which their reproductive benefits are maximized [32]. In the meantime, the choice of the host is important, which may contribute to the variation in the abundance of the fleas collected when the potential risk of flea-mediated diseases are assessed. In addition, the variation in the level of infection in host individuals is well known, indicating that some individuals may represent better patches for parasites than other individuals [20]. Namely, reservoirs providing better patches for parasites are usually defined as primary hosts.

Differences in flea infestation due to the age of the host may be influenced by differences in parasite aggregation. We tested the effect of age of small mammals on parasitic fleas and found that the probability of flea prevalence in adult hosts increased as the number decreased. This phenomenon has generally been linked to host immune responses and parasite-host associations. Firstly, it is known that immune defenses often deteriorate with increasing age [33], so that in smaller mammals the decline of antiparasitic defense has been reported with increasing age [34]. These hosts have different defenses that can affect the number of blood a flea can acquire [35]. Fleas were found to take more blood from juveniles and older animals than from sub-adult and adult animals [36]. The degree of deterioration of immune function with age may, however, be affected by environmental conditions (e.g. environmental stress) [37] and may also differ between males and females (due to faster aging of men) [38]. Secondly, differential parasite abundance in hosts belonging to different age cohorts has been reported for various host and parasite taxa [39–41]. However, the influence of host age on the distribution pattern of parasite abundance differs among different host-parasite associations. In some host-parasite associations, the abundance of parasites increases with the age of the host [39], while in other associations it increases or decreases in the youngest and oldest hosts compared to hosts of median age (called “adult hosts”)[40]. In fact, acquired resistance against parasites may be lower in young or old hosts than in median-age hosts. Young hosts may simply not have the time to develop parasite resistance, while older hosts may lose their ability to resist parasites due to immunosenescence [33, 42, 43]. As a result, hosts in the youngest and the oldest cohorts would have better habitat for parasites, so the relationship between parasite abundance and host age would be convex. If the negative effect of heavy parasite loads causes mortality mainly in young instead of old hosts, the abundance of parasites will increase with the age of the host. In all cases, fleas performed better in younger and older hosts than in middle-aged hosts.

In addition, the effect of the age of the host was strongly influenced by the effect of the sex of the host. In particular, from the point of view of resource acquisition (i.e. The size of the blood meal), there has been an improvement in the quality of cohorts of young and old among female, but not in male hosts, while, in the context of resource processing (digestion of blood), some trends in age-dependent host quality has been observed in male but not in female hosts [35]. In other words, the results of the Liberman study suggest that the age of the host does not predict unequivocally whether it is more or less beneficial for a flea [36].

Similarly, the sex of the host was also a determining factor influencing the abundance of fleas. Previous studies have shown the effect of host sex in flea reproduction [44,45]. In this study, we studied the consequences of host sex on flea abundance. Finally, we observed an increase in the number of flea infestations in small male mammals compared to females. The effect of host sex on parasite performance is essential to understanding the mechanisms of male-biased parasitism.

Under normal conditions, two non-mutually exclusive mechanisms are invoked to explain the variation in the number of fleas observed. Male hosts are characterized by higher mobility and poorer immunological defense than female. In general, fleas feed faster, take relatively more blood and digest more quickly when feeding on male rather than female, although the pattern of blood digestion related to the sex of the host depends on external conditions (relative humidity). In addition, fleas produced more eggs exploiting male hosts than female hosts [44]. This would lead to a better ability of parasites exploiting male rather than female because of the immunosuppressive effects of androgens, which would lead to fleas favoring male [45]. It is suggested that, in many cases, male hosts represent better patches for parasites than females. Thus, it can be seen that more flea prefers to stay in male hosts than in females. Consequently, a greater infestation of a male host than a female host in terms of abundance, prevalence and species richness of parasites, this has been reported for a wide variety of parasite and host taxa [46], although higher levels of parasite infection have been reported in some mammalian female [47].

The body weight of the hosts could be considered as an additional indicator of the number of fleas assembled. The results of this study showed that the number of fleas increased in small mammals weighing more than 59 grams. Hawlena [48] noted that the number of fleas, mainly *Xenopsylla cheopis* and *Xenopsylla Astia* (Siphonaptera: Pulicidae), caught in the wild (Rodentia: Muridae) increased with the weight of the host (and thus its age) at an optimum before decreasing. In practice, the body weight of the host increases with age. As mentioned above, the probability of flea infestation on the host increases with the age of the host. Therefore, the number of fleas would increase with the body weight of the host.

Associations between the aggregation of fleas and the abundance of their hosts vary according to different factors. This study confirmed that the variation in the number of fleas was due to different factors in the host.

It is concluded that the host is an important factor to consider when comparing parasite flea variation in natural rodent populations. It is necessary that the factors mentioned above be taken into account to control the abundance of parasitic fleas. Fitness related measures should be directly involved, such as reducing the number of small mammals and flea fleas, preventing and controlling flea-borne diseases.

## Conclusions

The results of the study show the distribution of small mammals and household parasitic fleas in the western province of Yunnan, China. In addition, a new discovery revealed by the hurdle binomial regression model is that there is a close relationship between the abundance of parasitic fleas and the characteristics of small mammals (e.g. species, age, sex, and body weight). Our study provides direct evidence to explore the relationship between parasitic fleas and the host by the characteristic of instinct and to reveal host structure plays an important role in the allocation of parasitic flea abundance in the ecosystem of the plague. The knowledge of parasitic flea factors will help control and predict the number of parasitic fleas in commensal rodent plague foci and further control the onset of the plague. Although it is relatively limited that the current model predicts the factors affecting the abundance of parasitic fleas. In addition to the characteristics of the host, the number of parasitic fleas may be affected by other potential factors (e.g. environment, climate, economy, human disturbance). Thus, the above factors should be measured and included in the regression model, it would be useful to establish benefits model to predict the abundance of parasitic fleas.

## Acknowledgment

The work was supported by the National Natural Science Foundation of China (No. 81460485; No. 81860565).

## References

1. Gallizzi K, Alloitteau O, Harrang E, Richner H. Fleas, parental care, and transgenerational effects on tick load in the great tit. Behav Ecol. 2008; 19(6):1225–34. https://doi.org/10.1093/beheco/arn083.

2. Gage KL. Factors affecting the spread and maintenance of plague. Adv Exp Med Biol. 2012; 954:79–94. https://doi.org/10.1007/978-1-4614-3561-7_11.

3. Yin JX, Dong XQ, Liang Y, Wang P, Siriarayaporn P, Thaikruea L. Human plague outbreak in two villages, Yunnan Province, China, 2005. Southeast Asian J Trop Med Public Health. 2007; 38(6):1115–9.

4. Cai WF, Zhang FX, Wang GL. Structure and community diversity of small mammals in plague natural focus of Yulong County and Gucheng District. Chin J Ctrol Endem Dis. 2015; 30:333–5.

5. Stenseth NC, Atshabar BB, Begon M, Belmain SR, Bertherat E, Carniel E, et al. Plague: past, present, and future. PLoS Medicine. 2008; 5(1):e3. https://doi.org/10.1371/journal.pmed.0050003.

6. Liang XC, Wang DS. Analysis of epidemic situation of Marmota himalayana plague natural focus in Gansu Provence. Bull Dis Ctrol Prev. 2011; 26(1):38–48.

7. Bai LQ, Wang JJ, Si XY. Overview of the Survelliance of Rodent Populations From 2001 to 2011 in Inner Mongolia. Inner Mongolia Med. 2014; 46(7):826–9.

8. Yin JX, Geater A, Chongsuvivatwong V, Dong XQ, Du CH, Zhong YH, et al. Predictors for presence and abundance of small mammals in households of villages endemic for commensal rodent plague in Yunnan Province, China. BMC ecology. 2008; 8:18. https://doi.org/10.1186/1472-6785-8-18.

9. Li JY, Zhao WH, Dong XQ, Liang Y, Hong M. Analysis on the current status of plague epidemics for Rattus flavipectus plague natural foci in Yunnan province. Chin J Endemiol. 2006; 25(6):654–7.

10. Otranto D, Dantas-Torres F, Napoli E, Solari Basano F, Deuster K, Pollmeier M, et al. Season-long control of flea and tick infestations in a population of cats in the Aeolian archipelago using a collar containing 10% imidacloprid and 4.5% flumethrin. Vet Parasitol. 2017; 248:80–3. https://doi.org/10.1016/j.vetpar.2017.10.023.

11. Lawrence AL, Hii S-F, Jirsová D, Panáková L, Ionică AM, Gilchrist K, et al. Integrated morphological and molecular identification of cat fleas (Ctenocephalides felis) and dog fleas (Ctenocephalides canis) vectoring Rickettsia felis in central Europe. Vet Parasitol. 2015; 210(3/4):215–23. https://doi.org/10.1016/j.vetpar.2015.03.029.

12. Krasnov BR. Functional and Evolutionary Ecology of Fleas:a Model for Ecological Parasitology Cambridge University Press: Cambridge, UK; 2008.

13. Giorgi MS, Arlettaz R, Christe P, Vogel P. The energetic grooming costs imposed by a parasitic mite (Spinturnix myoti) upon its bat host (Myotis myotis). Proc Biol Sci. 2001; 268(1480):2071–5. https://doi.org/10.1098/rspb.2001.1686.

14. Sánchez S, Gómez MS. Xenopsylla spp. (Siphonaptera: Pulicidae) in murid rodents from the Canary Islands: An update. Parasite. 2012; 19(4):423–6. https://doi.org/10.1051/parasite/2012194423.

15. Rouault E, Lecoeur H, Meriem AB, Minoprio P, Goyard S, Lang T. Imaging visceral leishmaniasis in real time with golden hamster model: Monitoring the parasite burden and hamster transcripts to further characterize the immunological responses of the host. Parasitol Int. 2017; 66(1):933–9. https://doi.org/10.1016/j.parint.2016.10.020.

16. Schmid-Hempe P, Ebert D. On the evolutionary of specific immune deference. Trends Ecol Evol. 2003; 18:27–32.

17. Khokhlova IS, Spinu M, Krasnov BR, Degen AA. Immune response to fleas in a wild desert rodent: effect of parasite species, parasite burden, sex of host and host parasitological experience. J Exp Biol. 2004; 207(16):2725–33. https://doi.org/10.1242/jeb.01090.

18. Lorange EA, Race BL, Sebbane F, Hinnebusch BJ. Poor vector competence of fleas and the evolution of hypervirulence in Yersinia pestis. J Infect Dis. 2005; 191(11):1907–12. https://doi.org/10.1086/429931.

19. Eisen RJ, Bearden SW, Wilder AP, Montenieri JA, Antolin MF, Gage KL. Early-phase transmission of Yersinia pestis by unblocked fleas as a mechanism explaining rapidly spreading plague epizootics. Proc Natl Acad Sci U S A. 2006; 103(42):15380–5. https://doi.org/10.1073/pnas.0606831103.

20. Poulin R. Evolutionary Ecology of Parasites: From Individuals to Communities. Princeton University Press: Princeton,U S A.; 2007.

21. Poulin R. Relative infection levels and taxonomic distances among the host species used by a parasite: insights into parasite specialization. Parasitology. 2005; 130(1):109–15.

22. Feng JM, Xu CD. Geographical distribution patterns of zonal plant community species diversity in Western Yunnan, China. Chin J Ecol. 2009; 28(4):595–600.

23. Zeileis A, Kleiber C, Jackman S. Regression Models for Count Data in R. J Stat Softw. 2008; 27(8):1–25.

24. Baughman AL. Mixture model framework facilitates understanding of zero-inflated and hurdle models for count data. J Biopharm Stat. 2007; 17(5):943–6. https://doi.org/10.1080/10543400701514098.

25. Yin JX, Dong XQ. Application of Hurdle Model in Identifying Predictors for Flea Abundance on Rats. Endem Dis Bull. 2010; 25(5):1–4.

26. Yin JX, Geater A, Chongsuvivatwong V, Dong XQ, Du CH, Zhong YH. Predictors for abundance of host flea and floor flea in households of villages with endemic commensal rodent plague, Yunnan Province, China. PLoS Negl Trop Dis. 2011; 5(3):e997. https://doi.org/10.1371/journal.pntd.0000997.

27. Yin JX, Zhong YH, Du CH, Dong XQ, Yang SH. Predictors for abundance of Rattus tanezumi in households of commensal rodent plague foci. Zhonghua liuxingbingxue zazhi. 2013; 34(2):157–9.

28. Wu AG, Li TY, Feng JM, Dong XQ. Study on the epidemiological significance related to community-structural difference of the rat plague host and vectors in Western Yunnan, China. Zhonghua liuxingbingxue zazhi. 2008; 29(4):346–50.

29. Krasnov BR, Sarfati M, Arakelyan MS, Khokhlova IS, Burdelova NV, Degen AA. Host specificity and foraging efficiency in blood-sucking parasite: feeding patterns of the flea Parapulex chephrenis on two species of desert rodents. Parasitol Res. 2003; 90(5):393–9. https://doi.org/10.1007/s00436-003-0873-y.

30. Krasnov BR, Shenbrot GI, Khokhlova IS, Poulin R. Relationships between parasite abundance and the taxonomic distance among a parasite’s host species: an example with fleas parasitic on small mammals. Int J Parasitol. 2004; 34(11):1289–97. https://doi.org/10.1016/j.ijpara.2004.08.003.

31. Khokhlova IS, Fielden LJ, Degen AA, Krasnov BR. Digesting blood of an auxiliary host in fleas: effect of phylogenetic distance from a principal host. J Exp Biol. 2012; 215(8):1259–65. https://doi.org/10.1242/jeb.066878.

32. Krasnov BR, Khokhlova IS, Shenbrot GI. Density-dependent host selection in ectoparasites: an application of isodar theory to fleas parasitizing rodents. Oecologia. 2003; 134(3):365–72. https://doi.org/10.1007/s00442-002-1122-2.

33. Gruver AL, Hudson LL, Sempowski GD. Immunosenescence of ageing. J Pathol. 2007; 211(2):144–56. https://doi.org/10.1002/path.2104.

34. Body G, Ferte H, Gaillard JM, Delorme D, Klein F, Gilot-Fromont E. Population density and phenotypic attributes influence the level of nematode parasitism in roe deer. Oecologia. 2011; 167(3):635–46. https://doi.org/10.1007/s00442-011-2018-9.

35. Liberman V, Khokhlova IS, Degen AA, Krasnov BR. Reproductive consequences of host age in a desert flea. Parasitology. 2013; 140(4):461–70. https://doi.org/10.1017/s0031182012001904.

36. Liberman V, Khokhlova IS, Degen AA, Krasnov BR. The effect of host age on feeding performance of fleas. Parasitology. 2011; 138(9):1154–63. https://doi.org/10.1017/s0031182011000758.

37. Hayward AD, Wilson AJ, Pilkington JG, Pemberton JM, Kruuk LE. Ageing in a variable habitat: environmental stress affects senescence in parasite resistance in St Kilda Soay sheep. Proc Biol Sci. 2009; 276(1672):3477–85. https://doi.org/10.1098/rspb.2009.0906.

38. Clutton-Brock TH, Isvaran K. Sex differences in ageing in natural populations of vertebrates. Proc Biol Sci. 2007; 274(1629):3097–104. https://doi.org/10.1098/rspb.2007.1138.

39. Fichet-Calvet E, Wang J, Jomaa I, Ben Ismail R, Ashford RW. Patterns of the tapeworm Raillietina trapezoides infection in the fat sand rat Psammomys obesus in Tunisia: season, climatic conditions, host age and crowding effects. Parasitology. 2003; 126(5):481–92.

40. Krasnov BR, Stanko M, Morand S. Age-dependent flea (Siphonaptera) parasitism in rodents: a host’s life history matters. J Parasitol. 2006; 92(2):242–8. https://doi.org/10.1645/ge-637r1.1.

41. Alarcos AJ, Timi JT. Parasite communities in three sympatric flounder species (Pleuronectiformes: Paralichthyidae): similar ecological filters driving toward repeatable assemblages. Parasitol Res. 2012; 110(6):2155–66. https://doi.org/10.1007/s00436-011-2741-5.

42. Johansen CE, Lydersen C, Aspholm PE, Haug T, Kovacs KM. Helminth parasites in ringed seals (Pusa hispida) from Svalbard, Norway with special emphasis on nematodes: variation with age, sex, diet, and location of host. J Parasitol. 2010; 96(5):946–53. https://doi.org/10.1645/ge-1685.1.

43. Praet N, Speybroeck N, Rodriguez-Hidalgo R, Benitez-Ortiz W, Berkvens D, Brandt J, et al. Age-related infection and transmission patterns of human cysticercosis. Int J Parasitol. 2010; 40(1):85–90. https://doi.org/10.1016/j.ijpara.2009.07.007.

44. Khokhlova IS, Serobyan V, Degen AA, Krasnov BR. Host gender and offspring quality in a flea parasitic on a rodent. J Exp Biol. 2010; 213(19):3299–304. https://doi.org/10.1242/jeb.046565.

45. Khokhlova IS, Serobyan V, Krasnov BR, Degen AA. Is the feeding and reproductive performance of the flea, Xenopsylla ramesis, affected by the gender of its rodent host, Meriones crassus? J Exp Biol. 2009; 212(10):1429–35. https://doi.org/10.1242/jeb.029389.

46. Patterson JE, Neuhaus P, Kutz SJ, Ruckstuhl KE. Patterns of ectoparasitism in North American red squirrels (Tamiasciurus hudsonicus): Sex-biases, seasonality, age, and effects on male body condition. Int J Parasitol Parasites Wildl. 2015; 4(3):301–6. https://doi.org/10.1016/j.ijppaw.2015.05.002.

47. Krasnov BR, Morand S, Hawlena H, Khokhlova IS, Shenbrot GI. Sex-biased parasitism, seasonality and sexual size dimorphism in desert rodents. Oecologia. 2005; 146(2):209–17. https://doi.org/10.1007/s00442-005-0189-y.

48. Hawlena H, Khokhlova IS, Abramsky Z, Krasnov BR. Age, intensity of infestation by flea parasites and body mass loss in a rodent host. Parasitology. 2006; 133(2):187–93. https://doi.org/10.1017/s0031182006000308.

